# Population structure in the MHC region

**DOI:** 10.1101/2021.10.25.465726

**Authors:** André Silva Maróstica, Kelly Nunes, Erick C. Castelli, Nayane S. B. Silva, Bruce S. Weir, Jérôme Goudet, Diogo Meyer

## Abstract

In his 1972 “The apportionment of human diversity”, Lewontin showed that, when averaged over loci, genetic diversity is predominantly attributable to differences among individuals within populations. However, selection on specific genes and genomic regions can alter the apportionment of diversity. We examine genetic diversity at the HLA loci, located within the MHC region. HLA genes code for proteins that are critical to adaptive immunity and are well-documented targets of balancing selection. The SNPs within HLA genes show strong signatures of balancing selection on large timescales and are broadly shared among populations, with low F_ST_ values. However, when we analyze haplotypes defined by these SNPs (i.e., which define “HLA alleles”), we find marked differences in frequencies between geographic regions. These differences are not reflected in the F_ST_ values because of the extreme polymorphism at HLA loci, illustrating challenges in interpreting F_ST_. Differences in the frequency of HLA alleles among geographic regions are relevant to bone-marrow transplantation, which requires genetic identity at HLA loci between patient and donor. We explore the case of Brazil’s bone-marrow registry, where a deficit of enrolled volunteers with African ancestry reduces the chance of finding donors for individuals with an MHC region of African ancestry.

## Introduction

In “The apportionment of human genetic diversity”, Richard Lewontin addressed a well-defined and answerable question: “how much of human diversity between populations is accounted for by more or less conventional racial classification?” With the genetic data available at the time and drawing on existing classifications of human races, he reached the unequivocal result that individuals assigned to different races are, on average, only slightly more genetically different than those from the same race.

Lewontin’s work redefined our way of thinking about, and referring to, human genetic variation, bringing an awareness that racial labels are arbitrary from the biological perspective, and largely socially constructed. Appropriately, subsequent studies shifted the focus to an understanding of how human genetic variation is distributed across the globe, what are the geographic scales at which such variation is observed (Rosenberg et al. 2005), and which evolutionary processes account for the observed patterns (Manica, Prugnolle, and Balloux 2005; Prugnolle, Manica, and Balloux 2005). What Lewontin referred to as “apportionment” has largely been recast in terms of “population structure”, and his approach to describing variation is now explored using metrics related to population genetic models, as is the case of studies using F_ST_.

Lewontin was aware that his main result referred to an “average” behavior over loci, and that variation in population structure among loci arises as a consequence of evolutionary sampling, as well as the locus-specific effects of natural selection. He explored this idea in (Lewontin and Krakauer 1973) by using the properties of the observed distribution of F_ST_ over loci to make inferences about natural selection. While this effort had limited success due to the difficulty in defining an appropriate null expectation for the distribution of F_ST_ (Nei and Maruyama 1975), the strategy has since become a central approach to study natural selection. In the genomic era, outlier approaches can be used (i.e., the detection of loci with extreme F_ST_), and the sheer scale of the data also allows parameter-rich models to be used to produce simulations that provide a null model (Nielsen 2005).

Here, we discuss the apportionment of genetic diversity for a particular set of loci, the HLA genes, which are located on chromosome 6 in a region of approximately 4 megabase known as the Major Histocompatibility Complex (MHC). The HLA genes have long attracted the interest of evolutionary biologists because of their unusually high levels of polymorphism (Bodmer 1972). The function of the proteins coded by HLA genes helps understand this unusual diversity: HLA proteins bind peptides and present them on the cell surface to T-cell receptors, triggering a cellular or humoral immune response if the MHC-peptide complex is recognized as foreign, as is the case for peptides from pathogens (Bodmer 1972; Klein and Sato 2000). The repertoire of peptides an individual can present on the cell surface depends on the HLA alleles which are present. As a consequence, pathogen diversity engages in a coevolutionary interaction with HLA loci. This interaction explains why HLA genes stand out as having the strongest evidence of balancing selection in the genome (Meyer et al. 2018), with (a) elevated non-synonymous substitution rates (Hughes and Nei 1992), (b) alleles with frequencies that deviate from neutral expectations (Hedrick and Thomson 1983), (c) DNA sequences with an excess of common variants (Bitarello et al. 2018), and (d) far more polymorphisms that are shared among species than expected under neutrality (Leffler et al. 2013).

Here, we investigate how selection affects the apportionment of genetic variation at HLA loci. The population structure at HLA loci is informative about the mode and timescale of selection, and the prevalence of local adaptation (e.g., selection favoring variants in a specific region) versus geographically widespread selection (e.g., selection favoring polymorphism which is shared among distantly related populations). An understanding of population structure at HLA loci is also relevant to Hematopoietic Stem Cell Transplantation (HSCT, also known as bone marrow transplantation), an important curative procedure used in the treatment of various forms of cancer and hematological diseases, where cells capable of generating hematopoietic tissue are transferred from a donor to a recipient. Successful HSCT requires genetic identity at HLA loci between patient and donor, a condition which is related to how genetic variation is apportioned. We will explore the relevance of population structure at HLA loci to challenges in planning the recruitment of unrelated donors for HSCT in Brazil, a country with a highly admixed population, with large components of European, African and Native American ancestry (Salzano and Bortolini 2005; Souza et al. 2019).

## Data analysis

### The 1000 genomes data

For our reanalysis of population structure in the MHC region, we have chosen the data for 20 non-admixed populations from Phase 3 of the 1000 Genomes Dataset (hereafter 1000G) (1000 Genomes Project Consortium and Adam Auton, Lisa D Brooks, Richard M Durbin, Erik P Garrison, Hyun Min Kang, Jan O Korbel, Jonathan L Marchini, Shane McCarthy, Gil A McVean, Gonçalo R Abecasis 2015). The 1000G data allows us to compare variation of the MHC to that in other genomic regions, to make calls for nucleotide positions within HLA loci, and to estimate which HLA alleles an individual carries (as described later). By “HLA alleles” we refer to the phased combination of variants within a locus, an important unit of analysis since it is the HLA alleles that define immunological phenotypes which contribute to disease and adaptation. Although the sampling structure of the 1000G is much coarser than that used by Lewontin (1972), we will be able to compare genetic variation among four major geographic regions (Africa, Europe, South Asia, East Asia).

### HLA allele and SNP calls

Genotype calls within HLA loci for the 1000 Genomes data are biased towards higher frequencies of reference alleles, due to mapping bias (Brandt et al. 2015). We therefore made new calls by extracting reads that either map to the MHC region or are unmapped (using the 1000 Genomes release describe by (Byrska-Bishop et al. 2021) and then performing a locus-specific mapping using known HLA alleles as references (Castelli et al. 2018). Throughout the text, we refer both to analyses for SNPs within HLA genes (in our case, the *HLA-A*, -*B*, and -*C*) and the MHC region (the broader region within which these and other genes of immunological function are contained). We also analyze diversity for “HLA alleles”, the phased combination of SNPs that defines an unique HLA protein sequence. Given our interest in functional variation at the HLA loci, we will analyze diversity among HLA alleles that correspond to unique protein sequences.

### F_ST_ estimates

In addition to revisiting Lewontin’s approach to quantifying the apportionment of genetic variation, we estimate F_ST_ to describe population structure. Our analyses will use the framework of Weir and Goudet (2017), where population-specific F_ST_ is computed based on allelic sharing and interpreted as a measure of relative kinship.

For genotypic data, for population *i*, we define the population-specific F_ST_ metric as:

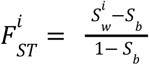

S refers to allele sharing between two individuals. For a bi-allelic locus, S assumes a value of one if the two individuals are homozygous for the same allele, zero if they are homozygous for different alleles, and 1/2 otherwise. Multiallelic loci can be accommodated in this framework by recoding them as a series biallelic markers. 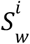 is sharing within population *i*, and *S_b_* is sharing between pairs of populations, averaged over pairs. Population-specific F_ST_ estimates the probability that two alleles drawn from population i are identical by descent (ibd) relative to a pair of alleles drawn from different populations. Averaging 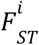 over populations provides the classically defined “overall F_ST_” (B. S. Weir and Clark Cockerham 1984). When estimating F_ST_ for a genomic region containing multiple SNPs, we use the “ratio of averages” approach, where the average value of the denominator and numerator are computed separately over SNPs, providing an unbiased estimator of F_ST_ (Ochoa and Storey 2021).

The F_ST_ estimates were computed with the Hierfstat R package (Goudet 2005). For the analysis of the entire chromosome, we filtered for biallelic SNPs and indels using VCFtools v.0.1.15 (Danecek et al. 2011) and converted from genotypic data to dosage format using Plink v.1.9 (Chang et al. 2015). In analyses of HLA alleles and SNPs within HLA loci, we used the genind2hierfstat and loci2genind functions (from the Hierfstat and Pegas packages) to format the data, and the fstat2dos and fs.dosage functions (from the Hierfstat package) to convert genotypic data to dosage and then calculate F_ST_.

### How is HLA diversity apportioned?

If Lewontin had HLA data available at the time of his study, what patterns would he have found? Lewontin’s approach consisted in comparing genetic diversity for progressively more inclusive groups (in his case, ranging from populations, to anthropologically defined races, to the entire species). He argued that when there are large differences between groups but little variability within them, measures of diversity for the species as a whole would be higher than those for each separate group. Using his approach, we find that heterozygosity at classical HLA class I loci (*HLA-A*, *HLA-B*, and *HLA-C*) is on average only slightly higher within regions or for the entire world (“total”), as compared to within populations (Table 1).

**Table 1.** Diversity at the HLA allele level and for the set of SNPs within HLA genes in the 1000 Genomes data, for *HLA-A*, -*B*, -*C*. Values refer to average heterozygosities over each form of grouping individuals (populations, geographic regions, total; H_p_, H_r_, H_t_, respectively). Total refers to the diversity present if all populations are treated as a single group. The regions (and populations) are: Africa (Gambian Mandinka, Mende, Esan, Yoruba, Luhya); Europe (Finnish, Iberian, Toscani, CEU - Northern Europeans, British); East Asia (Southern Han Chinese, Kinh Vietnamese, Japanese, Han Chinese, Dai Chinese); and South Asia (Bengali, Punjabi, Gujarati, Tamil, Telugu).

|  | HLA allele level Heterozygosity | SNP level Heterozygosity |
| --- | --- | --- |
| Populations | 0.904 | 0.170 |
| Regions | 0.916 | 0.172 |
| Total | 0.948 | 0.176 |

In Lewontin’s framework, the apportionment of HLA variation was computed as the relative gain in diversity obtained by amalgamating samples at higher level of grouping. Thus, the fraction of variation contributed by various levels of population structure can be computed for HLA allele level diversity as follows:

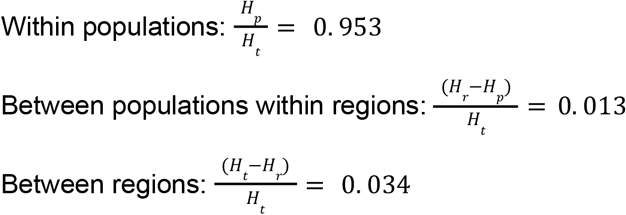

For SNP level heterozygosities, these same calculations result in 0.965, 0.012, 0.022 of the total variation being due to differences within populations, between populations within regions, and between regions, respectively.

Clearly, the vast majority of HLA variation, both at the HLA allele and SNP levels, lies within populations, and little additional variation is accrued by examining higher grouping. Notice that this does not imply that there are few differences between populations and geographic regions, but rather that they add very little diversity relative to that which is already present within populations. This overall pattern has been found in other studies of HLA variation across populations, which used hierarchical F_ST_ approaches, and also accounted for the level of divergence among HLA alleles ((Buhler and Sanchez-Mazas 2011; Sanchez-Mazas 2007; Meyer et al. 2006)).

While the overall picture of HLA variation is clearly one of high diversity within individual populations, analyses restricted to HLA genes are unable to address important questions. First, inferences about population structure of a specific locus requires placing it in a genome-wide context. This is difficult to accomplish when we use variation for entities such as “HLA genes” since the size of the locus, the density of SNPs, and the mutational mechanism all confound the interpretation of variation and, therefore, of the pattern of population structure. Secondly, natural selection may act heterogeneously across populations, so selective episodes that are specific to a population or region would be unnoticed in a survey of “overall” patterns of population structure, averaged over the entire dataset. This issue calls for approaches examining the evolutionary history of individual populations or lineages (Bruce S. Weir et al. 2005; B. S. Weir and Hill 2002; Bruce S. Weir and Goudet 2017), and defining a marker type that can be compared among genomic regions.

An objective way to compare the apportionment of variation at HLA genes with genome-wide values is to contrast estimates for SNPs within the MHC region, and within HLA genes, to genome-wide values (thus controlling for marker types) and using the same set of populations and individuals (thus controlling for population and individual sampling effects). In addition, using the population-specific FST approach, inferences about specific populations or regions (as opposed to overall values, averaged over all populations) can be made. Here, we use the 1000 Genomes data and the population-specific FST framework to revisit the question of how HLA diversity is apportioned by focusing on contrasts between SNPs in the MHC region, within HLA genes, and in the remainder of chromosome 6.

### MHC diversity in a genomewide context

To visualize how population structure in the MHC region compares to genomewide values, we estimated the population-specific F_ST_ for 5Mb windows along chromosome 6 (Figure 1). The five African populations have lower population-specific F_ST_ throughout the entirety of chromosome 6, reflecting higher diversity and thus lower within-population kinship among individuals, as compared to kinship within populations from other regions. However, within the MHC region Africans have a higher population-specific F_ST_ than in the remainder of the chromosome, and Asian and European populations show the opposite, i.e., a reduction in their FST values within the MHC (Figure 1, see MHC region delimited by vertical lines). The increase seen for F_ST_ in the African MHC region expresses the fact that, for this genomic region, Africans do not have a markedly higher diversity (and thus lower degree of within population kinship) than do other populations.

**Figure 1.**
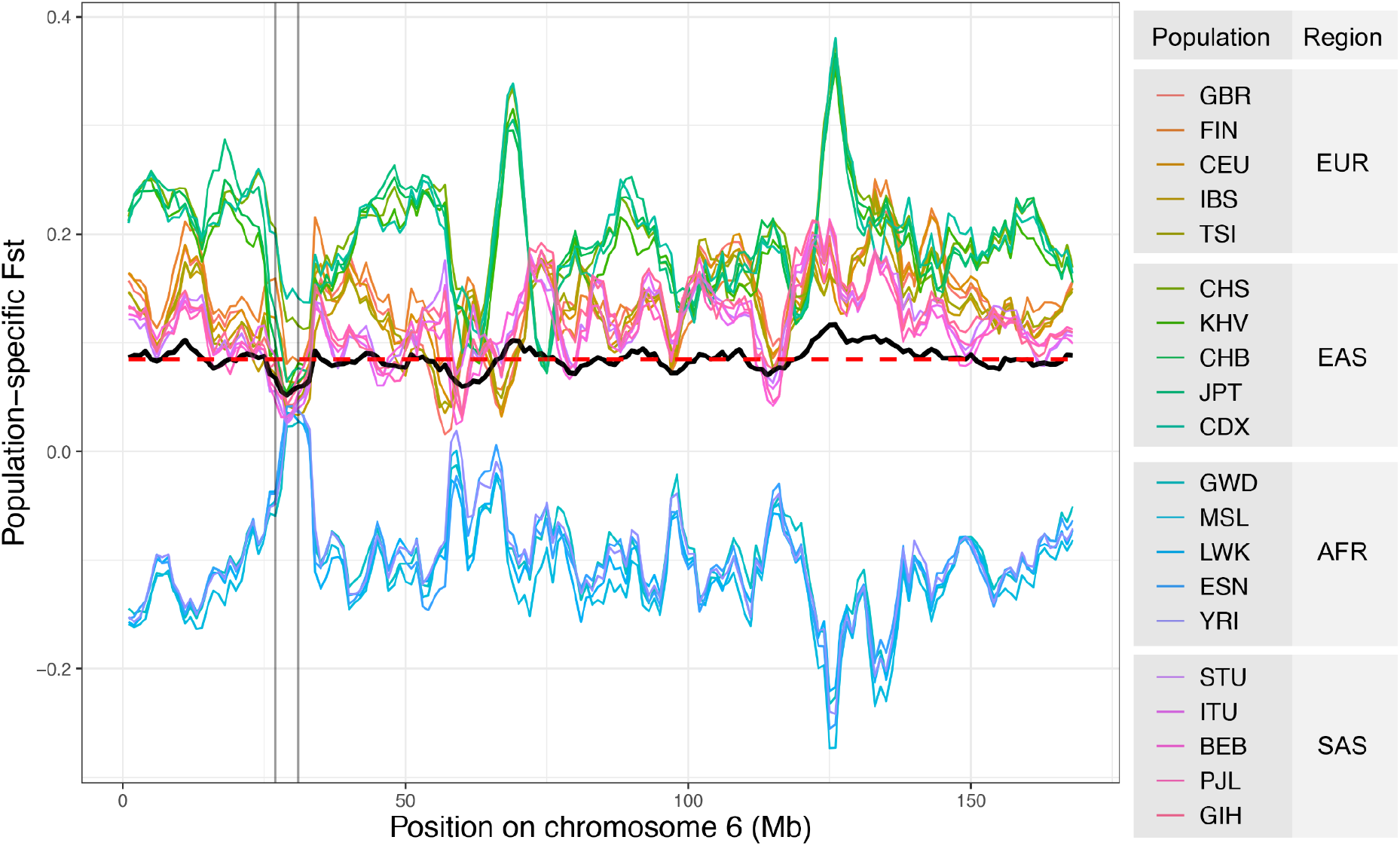
Population-specific F_ST_ across chromosome 6, in windows of 5Mb, and step-size of 1Mb. Each colored line represents a specific population; the black line represents the overall F_ST_, for each window; the red dashed line is the average overall F_ST_ for the entire chromosome. The vertical lines delimit the MHC region, within which HLA genes are contained. See supplementary Table S1 for 1000G abbreviations of population and region names.

The overall F_ST_ (i.e., the average of population-specific F_ST_ values), reaches its lowest value in chromosome 6 within the MHC (black line in Figure 1). The outlier status of the windows in the MHC region can be visualized by comparing the distribution of their F_ST_ values to that of windows in the remainder of chromosome 6 (Figure 2, top panel). The overall F_ST_ for the set of SNPs contained within the three HLA genes (a subset of the MHC region) is even more extreme (Figure 2A, vertical arrow). The apportionment of diversity among regions can also be computed by treating all the populations in a geographic region as a single group and computing an overall F_ST_ among regions (Figure 2, bottom panel). Once again, the F_ST_ values for SNPs within the MHC and HLA genes occupy an extreme position, indicating unusually low F_ST_ with respect to the remainder of chromosome 6.

**Figure 2.**
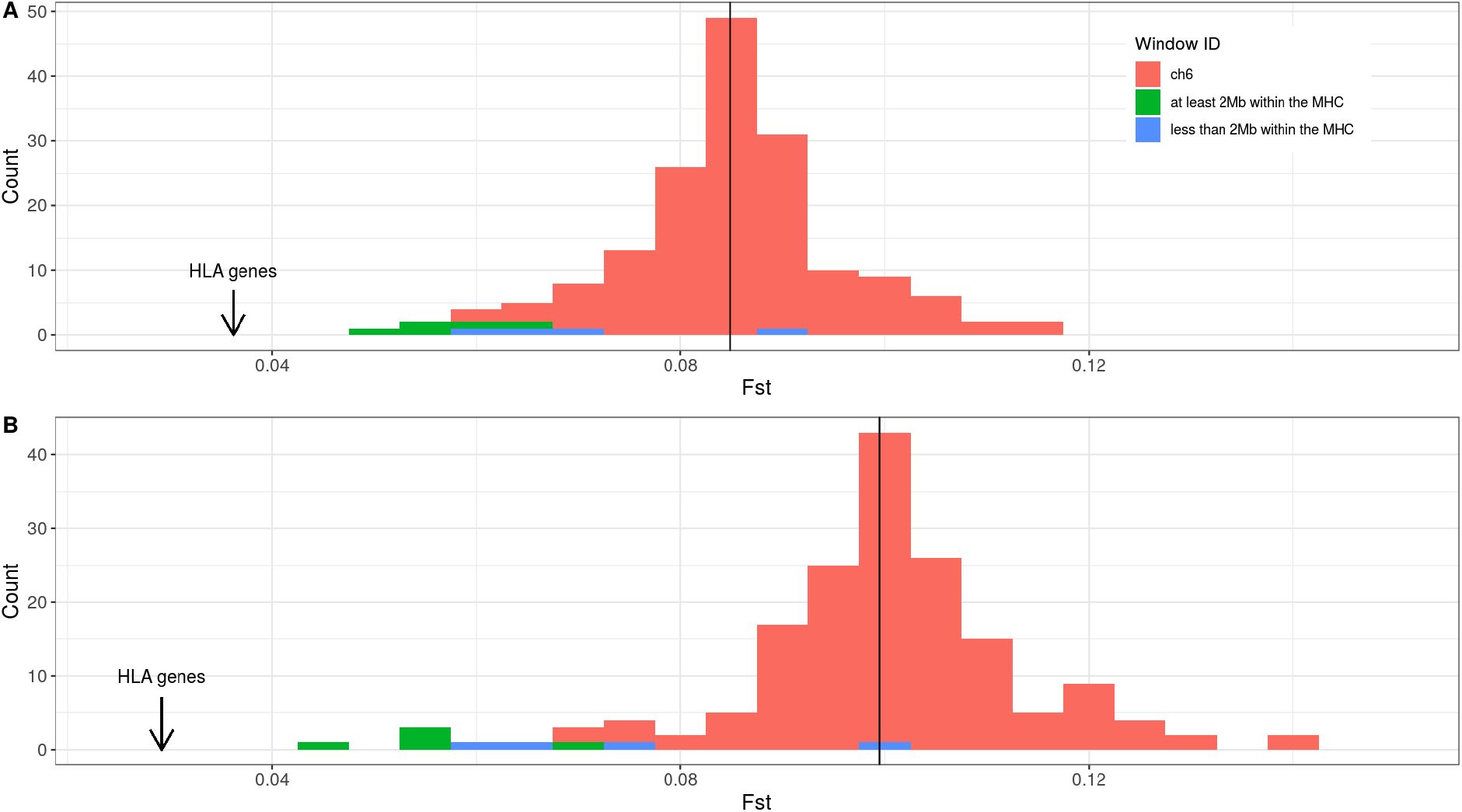
Distribution of overall F_ST_ for 168 windows along chromosome 6. The F_ST_ values for windows that have at least 2Mb contained within the MHC are shown in green, and the windows that intersect less than 2Mb of MHC are shown in blue, while the windows completely outside the MHC region are in red. The arrow indicates the overall F_ST_ for the SNPs contained within the three HLA loci we studied (HLA-A, -B, -C), and the vertical line is the average F_ST_ over all windows. (A) overall F_ST_ among populations. (B) overall F_ST_ among continents.

The outlier status of the MHC region SNPs, particularly those contained within HLA genes, supports a role for natural selection favoring the maintenance of diversity and resulting in extensive sharing of polymorphisms among regions and populations.

### Local adaptation at HLA loci

The overall F_ST_ for a locus captures the influence of evolutionary sampling and natural selection, averaged over populations. However, it is possible that selection acts in a population-specific manner, with the overall pattern masking signatures within specific populations (Bruce S. Weir et al. 2005). This has proved a relevant issue for studies of MHC diversity, which have recently found examples of specific populations with signatures of increased FST, in the opposite direction to the overall FST values presented above.

For a set of closely related African populations (African-American, Nigerians and Gambians), Bhatia and coworkers (Bhatia et al. 2011) found that the F_ST_ in the MHC region significantly exceeded that of the rest of the genome. For another set of African populations (Patin et al. 2017) also found an excess differentiation at the MHC region for a sample of Bantu speakers, relative to genomewide. In an analysis of Native Americans, (Nunes et al. 2021) used microsatellites within the MHC and spread throughout the genome to show that F_ST_ in the MHC exceeded neutral expectations (obtained by simulations). Finally, (Brandt et al. 2018) found that pairwise contrasts between East Asian populations from the 1000G showed unusually high F_ST_, when compared to genome-wide averages.

Local adaptation of HLA alleles is also supported by the study of admixed populations, with several studies documenting an excess of African ancestry within the MHC region (Norris et al. 2021; Meyer et al. 2018), consistent with the hypothesis that alleles of African ancestry were advantageous in the admixed populations. A similar signature was also found for the Native American component in admixed Mexicans (Zhou, Zhao, and Guan 2016), again indicating that a subset of locally advantageous alleles rose in frequency after the onset of admixture.

Thus, while there is strong evidence of reduced overall F_ST_ for SNPs in the MHC region and HLA genes (Figures 1 and 2), studies that queried admixed or closely related sets of populations identified instances where F_ST_ for the MHC region is in fact higher, and not lower, than genome-wide averages. These findings are consistent with selection favoring locally adapted variants, and emphasizes the importance of considering that selective regimes at HLA loci may differ, depending on the timescale being considered (Garrigan and Hedrick 2003).

### HLA diversity and transplantation

Patients suffering from hematological diseases, including forms of cancer, can be treated with stem cells harvested from the bone marrow of a donor in a process called Hematopoietic Stem Cell Transplantation (HSCT). The ideal setting for this procedure is for donor and recipient to match at 5 HLA loci (*HLA-A*, -*B*, -*C*, -*DRB1*, -*DQB1*, referred to as 10/10 matching), with “matching” defined by identity at the protein level for HLA alleles. However, because the expected heterozygosities at the HLA allele level are typically greater than 0.90 within populations (Table 1), the chance that two unrelated individuals will match at five loci is extremely low. Therefore, the first option for patients is to search for compatible donors among close relatives. However, fewer than 30% of patients find a compatible donor among relatives, and their next option is to seek donors in registries.

An important question for registries is whether the genetic ancestry of the patient and the donor influences the chance of a match. The population structure of HLA loci is potentially informative about this question. For the two most common ancestries in Brazil (European and African), we computed the population-specific and overall F_ST_ for populations in the 1000G, and found values to be very low (Figure 3, overall F_ST_ =0.03 at the HLA allele level). At first sight, this appears to indicate that there is an extensive sharing of alleles among Africans and Europeans. However, for extremely polymorphic loci, as is the case for HLA, low F_ST_ is possible even in the absence of extensive sharing of alleles among populations (Hedrick 2005; Balloux et al. 2000), making it misleading to associate low F_ST_ to lack of differentiation.

**Figure 3.**
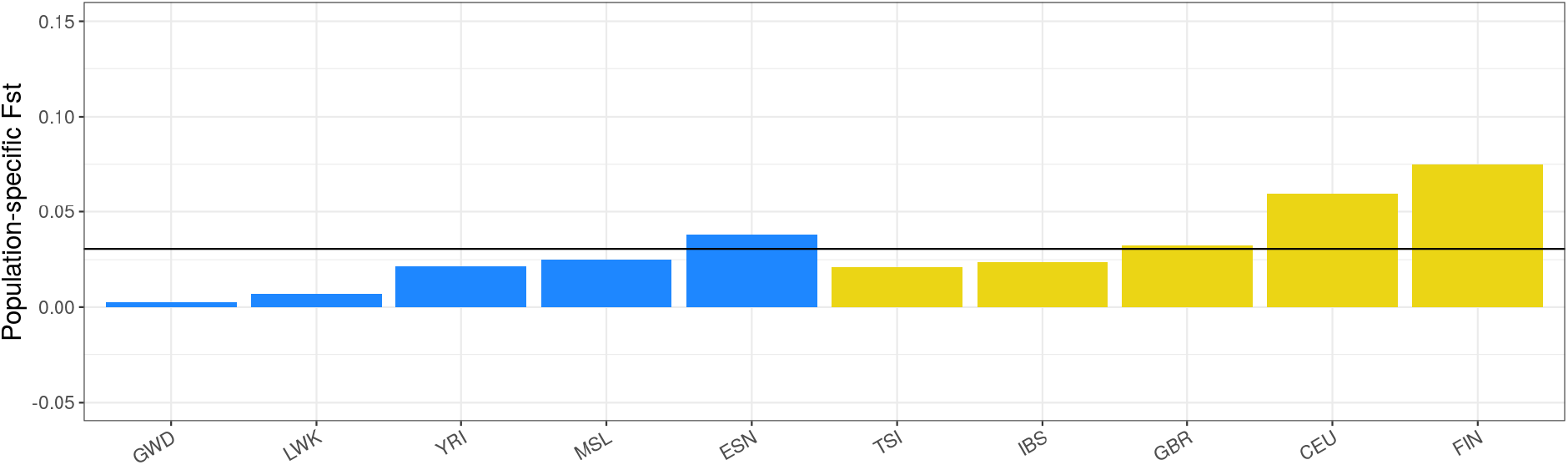
The population-specific F_ST_ values for African (in blue) and European (in yellow) populations in the 1000G, estimated for diversity at the level of HLA alleles for HLA-A, -B, -C. The horizontal line is the average over population-specific F_ST_, with a value of 0.03.

To visualize the differences in allele frequencies between African and European populations, we identified a set of HLA alleles that are either exclusive or at least three-fold more common in Africa, as compared to Europe, in the 1000G data. Figure 4 illustrates how 18 HLA-A and 20 -B alleles, which collectively contribute to approximately 60% cumulative frequency in Africa, reach less than 10% in Europe (for HLA-C alleles the differences among regions are smaller). The striking differences in allele frequencies among regions for HLA-A and -B are also observed in an independent dataset containing data from multiple populations (Solberg, O. D., Mack, S. J., Lancaster, A. K., Single, R. M., Tsai, Y., Sanchez-Mazas, A., & Thomson, G. 2008) Supplementary Figure S1-S3). This analysis shows how the distribution of HLA alleles is in fact geographically structured, but as a consequence of the extreme polymorphism, the overall F_ST_ is low (Hedrick 2005; Balloux et al. 2000). We do not see this as a limitation of F_ST_ (as do others, e.g., (Hedrick 2005; Jost 2008)), but as an expression of the information F_ST_ conveys, which is related to the evolutionary history of the population and locus of interest, rather than serving as a direct measure of differences in allele frequencies. Here, low F_ST_ reflects the high diversity (and low kinship) both within and between regions, a key feature of HLA polymorphism.

**Figure 4.**
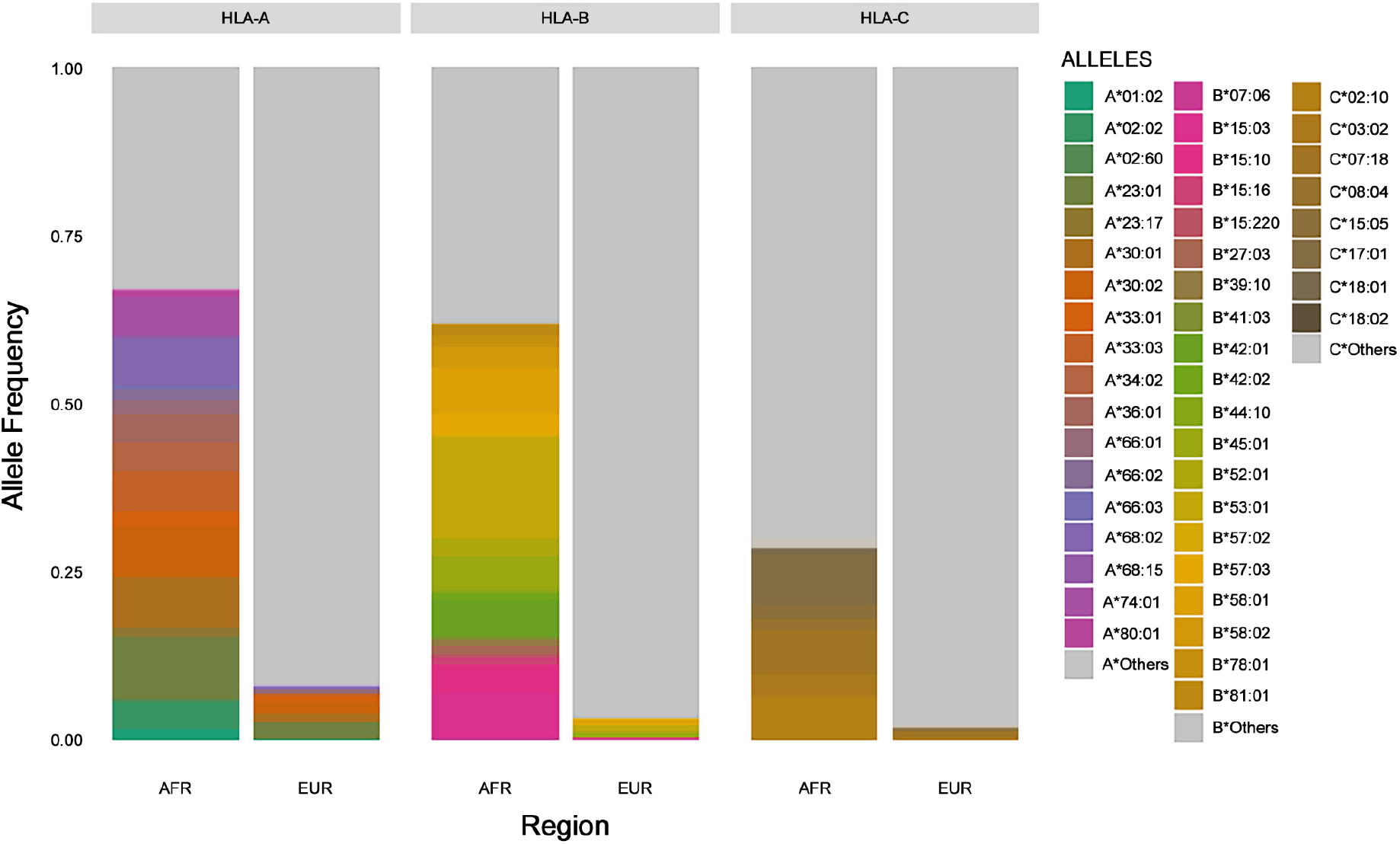
Frequencies of a subset of HLA alleles, for African and European populations (1000G). To visualize the differences in allele frequencies between regions, we selected alleles that were either exclusive to Africa, or found at frequencies that were at least threefold greater in Africa with respect to Europe. We identify each allele by a distinct color. “Others” (in gray) refers to the cumulative frequency of all alleles that are either exclusively found in Europe, or that do not occur threefold more frequently in Africa than Europe, or that are shared between them.

What are the consequences of the large differences in allele frequencies between geographic regions (Figure 4) to the chances of finding a matching donor in a repository? Brazil is a highly admixed country, with the largest population descended from Africans outside Africa (Ade Ajayi 1998), and a history of extensive migration from Europe. There is also an important but lower proportion of Native American ancestry, in particular in the north of the country. In Nunes et al (2020), we therefore quantified how an individual’s ancestry influences their chances of finding a matching donor in Brazil’s Redome, the Brazilian Bone Marrow Donor Registry. In a survey of over 8,000 admixed individuals, we showed that those who self-identify as “Black” have a 57% reduction in their chances of finding a matching donor, with respect to those who self-identify as “White”. When dividing the cohort into groups based on genomewide ancestry, there is a 60% reduction in chances of finding a donor in a group of 1589 individuals with the greatest African ancestry, with respect to those with the least African ancestry. Finally, when the question is recast with respect to genetic ancestry specific to the MHC region, we found that individuals with two MHC regions with African ancestry have a 75% reduction in chances of finding a donor (Figure 5A), with respect to those with two MHC regions identified as European.

**Figure 5.**
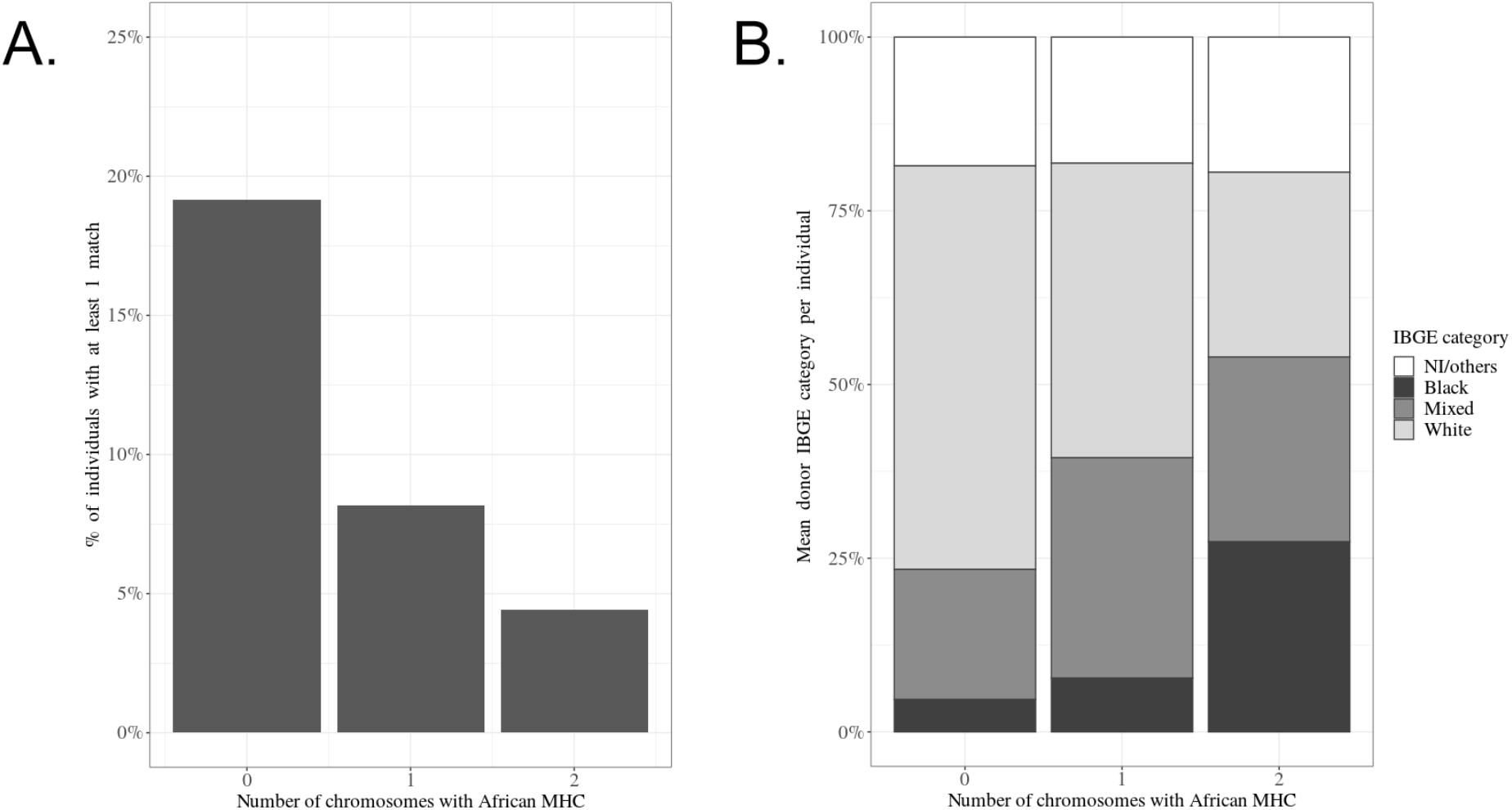
Ancestry and donor availability in the Brazilian bone marrow registry (for matching at both alleles at 5 loci, or 10/10 matching). Admixed individuals were classified regarding the genetic ancestry of their MHC region, using RFMIX (“RFMix: A Discriminative Modeling Approach for Rapid and Robust Local-Ancestry Inference” 2013). Each individual was assigned a value, referring to the number of chromosomes for which the MHC region was African: 0, 1, or 2. (A) Effect of African ancestry in the MHC on the proportion of individuals that find at least one match in the registry. The y axis represents the proportions of individuals who found at least one compatible donor. (B) The self-assigned identifier (a proxy coarsely related to ancestry) of potential donors, with respect to the genetic ancestry in the MHC region. Notice that for individuals with 2 African chromosomes in the MHC region, “Black” and “Mixed” make up the largest fraction of potential donors. (Adapted from Nunes et al., 2020).

The lower chances of finding a donor for individuals with African ancestry genomewide (and in particular for the MHC) is a consequence of REDOME’s underrepresentation of individuals of African ancestry, as documented in their statistics for self-identified labels used by the Brazilian Institute of Geography and Statistics (IBGE). Whereas the categories “Black” and “Mixed” correspond to 53.9% of Brazilians, they only make up 30.6% of registered donors in REDOME. Similar results were found for the National Marrow Donor Program (NMDP), where individuals who self-identify as “Black” have a 5-fold lower chance of finding a donor, compared to those who identify as “White” (Gragert et al. 2014).

Would increasing the proportion of individuals with African ancestry in the registries increase the chances of patients with a greater African ancestry in finding a compatible donor? Nunes et al (2020) found that individuals with a higher proportion of African ancestry in the MHC on average find proportionally more donors identified as “Black” or “Mixed” (Figure 5B). This suggests that for individuals with greater African ancestry, an increase in the proportion of donors with African ancestry will contribute to their chances of finding a compatible donor.

It may seem surprising to refer to “Black” and “Mixed” within a study that celebrates the very paper in which Lewontin provided a categorical rejection of racial categories. We believe reference to these categories is necessary in the context of the questions we address here, since government institutions routinely still employ these categories. It is appropriate to investigate their bearing on issues of public health, including the possible outcome of refuting their utility. Our results show that, in the case of finding matching donors, it is ultimately the genetic ancestry of the MHC which most strongly defines the chances of finding a match.

## Conclusion

The analysis of population structure at HLA connects with themes that Richard Lewontin explored in his scientific work and his communication about science to the non-specialized audience.

Lewontin’s writings on evolutionary biology recurrently emphasized the challenge of delimiting units of analysis, both at the morphological and genetic levels (Lewontin 2000)Lewontin, 2000). Indeed, there is no single correct way to define genotypes for the population structure analyses. We have shown that results for the diversity of SNPs and HLA alleles are both informative, but convey different types of information. The SNP level analysis captures the fact that most nucleotide level variation is widely shared among populations and regions (as in fact they are shared among species, (Leffler et al. 2013). However, such extensive sharing at the SNP level does not translate to sharing of HLA alleles among populations and regions, with the repertoire of HLA alleles differing markedly among regions (Figure 4). The decision with respect to which definition of “HLA genotype” will be used has a bearing on the distribution of genetic variation.

Lewontin’s work also emphasized the context-dependency of evolutionary trajectories, with empirical and analytical explorations of fitness surfaces that arise when interactions among loci and changing environments are considered (Lewontin and White 1960). The study of HLA genes provides an example of how a locus can carry the signatures of distinct selective regimes, at different timescales: strong evidence of balancing selection at the level of SNPs, when long timescales are considered, and evidence of recent selection, favoring locally adapted HLA alleles, at recent timescale (involving divergence between populations inhabiting the same continent, or recently admixed populations).

Finally, Lewontin was explicit about his political views and assumed that scientists’ technical work reflected their inevitable (although not always stated) political perspectives. The study of HLA population structure reveals how a technical question, regarding the degree of population structure in a genomic region, can have a bearing on public health issues in admixed populations. We ourselves were stimulated to investigate the effects of ancestry on the chances of finding donors by both recent scientific work in the United States (Gragert et al. 2014),and by websites maintained by patient-driven organizations (for example, https://blackbonemarrow.com/ and https://bonemarrowwish.org/). In the case of Brazil, the long history of slavery lies at the root of a pattern of systemic racism (Ribeiro 2019), and has resulted in a social organization where individuals of African ancestry have reduced access to quality healthcare, and are overburdened by many diseases, including COVID-19 (Martins-Filho et al. 2021). This calls for, among other efforts, research with the potential to guide institutional health care strategies. In the specific case of the Brazilian registry, strategies to recruit donors of African ancestry are a direct recommendation of the genetic analyses.

## Acknowledgements

We thank the organizers of this special issue for their invitation to contribute. DM thanks the members of the Evolutionary Genetics Laboratory at the University of São Paulo for helpful discussions.

## Authors’ contributions

Conceptualization (DM)

Data curation (KN, ASM, EC, NSBS)

Formal analysis (KN, ASM, DM, JG, BSW, NSBS, EC)

Funding acquisition (BSW, DM)

Investigation (KN, ASM, DM, BSW, JG)

Methodology (EC)

Supervision (DM, JG, BSW)

Writing - original draft (DM)

Writing - review & editing (DM with input from all authors)

