## Supplemental information for "Population structure in the MHC region"

1. Universidade de São Paulo, Departamento de Genética e Biologia Evolutiva, São Paulo, SP, Brazil.

2. Universidade Estadual Paulista, Faculdade de Medicina de Botucatu, Departamento de Patologia, Botucatu, SP, Brazil.

3. Department of Biostatistics, University of Washington, Seattle, Washington 98195.

4. Department of Ecology and Evolution,Swiss Institute of Bioinformatics, University of Lausanne, 1015 Switzerland.

**Supplementary Table S1.** Description of 1000G populations and geographic regions and sample size evaluated in this study.

| Population name | Abbreviations of population names | Geographic Region | Abbreviations of geographic region names | Number of samples |
|---|---|---|---|---|
| Esan | ESN | African | AFR | 96 |
| Gambian Mandinka | GWD | African | AFR | 107 |
| Luhya | LWK | African | AFR | 99 |
| Mende | MSL | African | AFR | 85 |
| Yoruba | YRI | African | AFR | 104 |
| British | GBR | Europe | EUR | 90 |
| CEPH | CEU | Europe | EUR | 99 |
| Finnish | FIN | Europe | EUR | 98 |
| Iberian | IBS | Europe | EUR | 107 |
| Toscani | TSI | Europe | EUR | 107 |
| Bengali | BEB | South Asian | SAS | 86 |
| Gujarati | GIH | South Asian | SAS | 101 |
| Punjabi | PJL | South Asian | SAS | 94 |
| Tamil | STU | South Asian | SAS | 98 |
| Telugu | ITU | South Asian | SAS | 101 |
| Dai Chinese | CDX | East Asian | EAS | 93 |
| Han Chinese | CHB | East Asian | EAS | 102 |
| Southern Han Chinese | CHS | East Asian | EAS | 105 |
| Japanese | JPT | East Asian | EAS | 103 |
| Kinh Vietnamese | KHV | East Asian | EAS | 99 |

**Supplementary Figure S1.** Frequencies and geographic distribution of a subset of HLA-A alleles, according to the compilation by Solberg et al., (2008). The subset of HLA alleles were selected using the criteria of being exclusively in the African populations, or found at frequencies that were at least threefold greater in African with respect to European population in the 1000 Genome data (1000G). The frequency distribution maps were obtained from the website (http://www.uvm.edu/~igdawg/software/browser-beta.html).

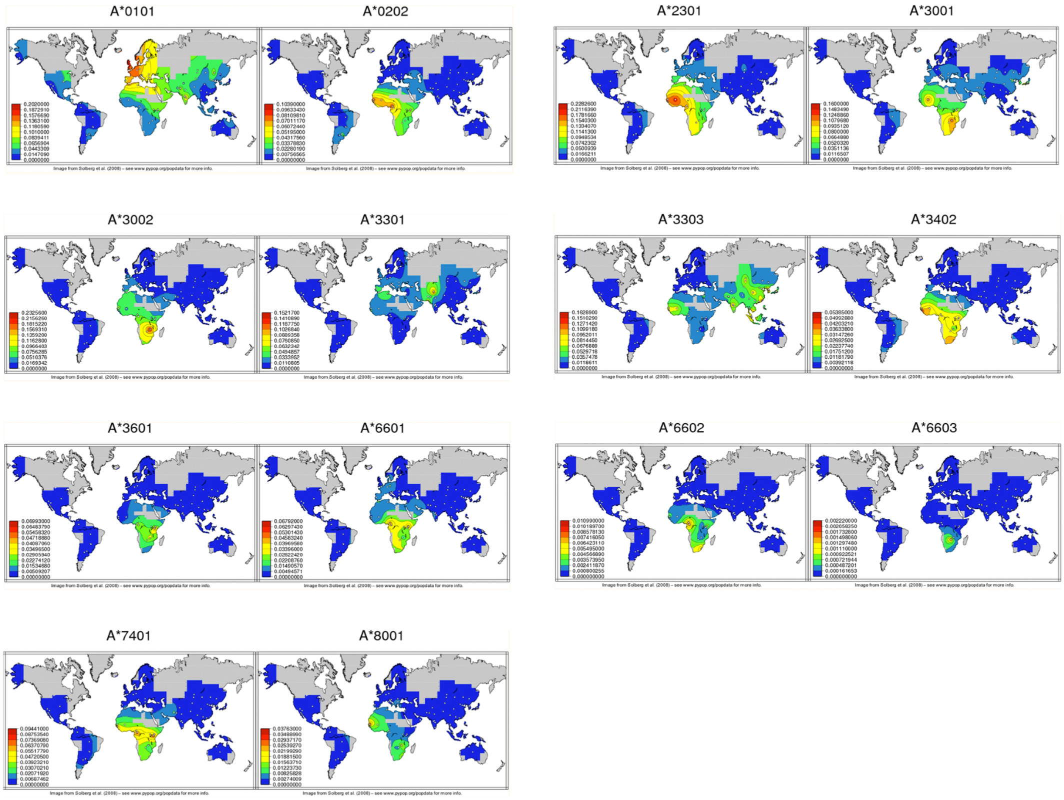

**Supplementary Figure S2.** Frequencies and geographic distribution of a subset of HLA-B alleles, according to the Solberg et al., (2008). The subset of HLA alleles were selected using the criteria of being exclusively found in the African populations, or found at frequencies that were at least threefold greater in Africans with respect to Europeans in the 1000 Genome data (1000G). The frequency distribution maps were obtained from the website (http://www.uvm.edu/~igdawg/software/browser-beta.html).

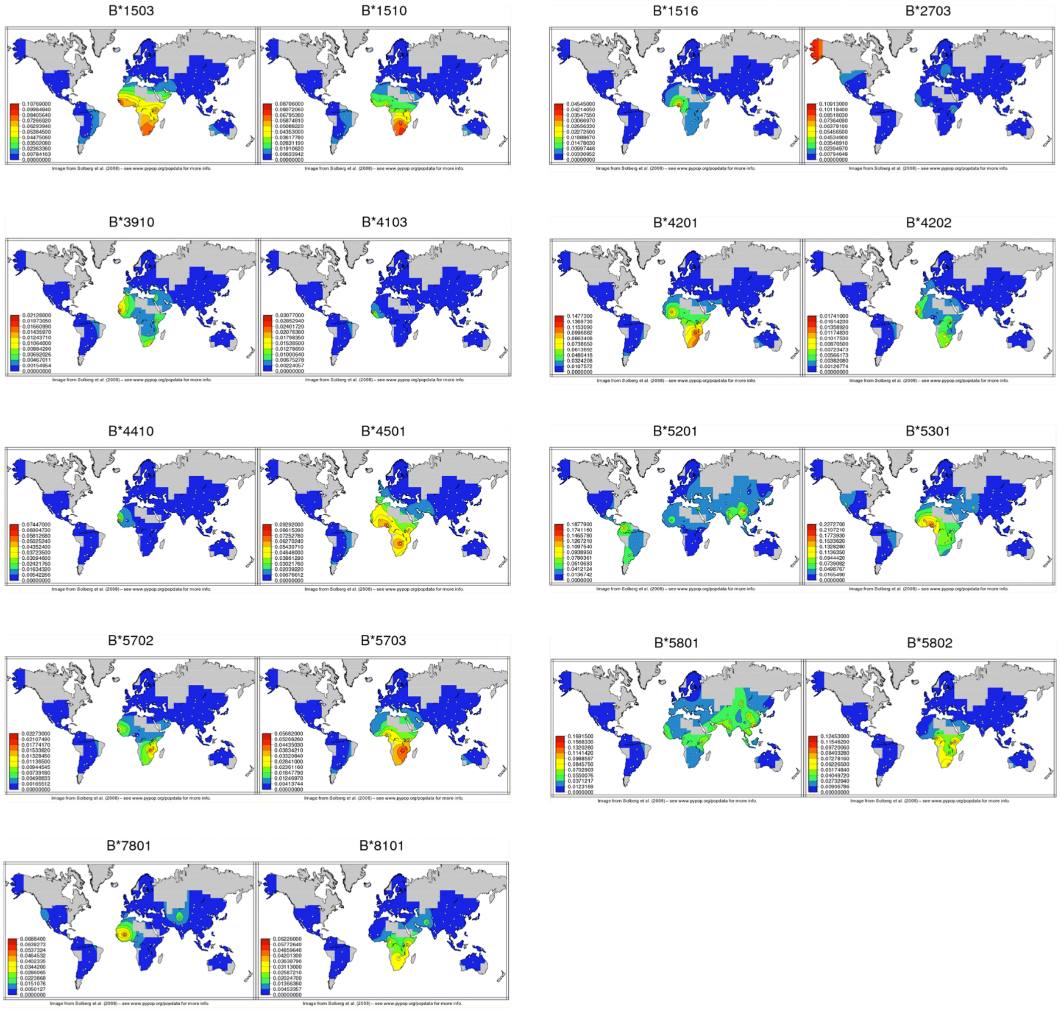

**Supplementary Figure S3.** Frequencies and geographic distribution of a subset of HLA-C alleles, according to the Solberg database compilation (Solberg et al., 2008). The subset of HLA alleles were selected using the criteria of exclusively in the African population, or found at frequencies that were at least threefold greater in African with respect to European population from 1000 Genome data (1000G). The frequency distribution maps were obtained from the website (http://www.uvm.edu/~igdawg/software/browser-beta.html).

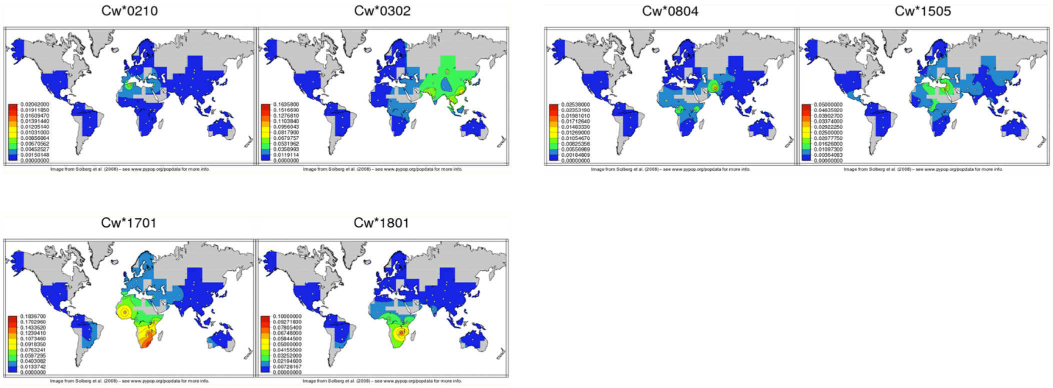
